# PPi plays a role in sugar starvation tolerance

**DOI:** 10.1101/2021.01.21.427394

**Authors:** Seedahmed A. Mohammed

**Affiliations:** Department of Biology and Biotechnology, Faculty of Science and Technology, AlNeelain University, Khartoum, Sudan

## Abstract

H^+^-PPase acidifies the endo-membrane compartments utilizing the energy of PPi hydrolysis. This acidification creates an electrochemical proton gradient that powers the secondary active transport and allows for vacuolar accumulation of several materials against their concentration gradients. The hydrolysis of the cytosolic PPi is necessary for the forwardness of the PPi-generating reactions. However, information is lacking on the role of PPi in adaptation to sugar starvation and low energy status in plants. Here, several mutants lacking the functional H^+^-PPases were used to illuminate this role. Three alleles of *fugu5* mutants defective in type I H^+^-PPase exhibited better tolerance to sugar starvation than wild-type plants, when grown on ½ and full-strength MS media under photosynthesis-constraining low light intensity. The PPi level in *fugu5* mutants was significantly higher than its level in wild type and type II H^+^-PPase-defective mutants. SnRK1 (Sucrose-non-fermenting1-Related kinase-1) plays a central role in the coordination of the plant transcriptome to the energy signals. SnRK1 senses the energy depletion in plant cells, and controls the expression of genes and phosphorylation of proteins in a way that promotes catabolism, and inhibits anabolism. Sugar starvation significantly induced the SnRK1 phosphorylation activity in wild type and type II H^+^-PPase lacking mutants. Whereas the activity remained unchanged in sugar-starved *fugu5* mutants. This is possibly achieved through activation of PPi-dependent enzymes. Results suggest that the high PPi level in *fugu5* mutants might contribute to more efficiently use of low level of ATP under sugar starvation and low light conditions.

## Introduction

The proton-pumping inorganic pyrophosphatases (H^+^-PPases) acidify the endomembrane compartments using the energy stored in the phosphoanhydride bond of pyrophosphate to drive protons across endomembranes (Maeshima, 2001; Gaxiola et al., 2007). Two types of H^+^-PPases exist in plant cell. They differ in primary structures, ions sensitivity and localization. Type I H^+^-PPase that localizes to the tonoplast is dependent on K^+^ for its activities and moderately sensitive to the inhibition by Ca^2+^, whereas, type II H^+^-PPase that localizes to golgi and trans-golgi networks is K^+^ insensitive but extremely Ca^2+^ sensitive (Drozdowicz et al., 2000).

In Arabidopsis genome there is a single type I H^+^-PPase-encoding gene, namely, *AtVHP1;1, FUGU5,* or *AVP1;* and two type II H^+^-PPase-encoding genes, namely, *AtVHP2;1,* or *AVP2;1,* and *AtVHP2;2,* or *AVP2;2* (Drozdowicz & Rea, 2001; Segami et al., 2010). The cellular content of type II H^+^-PPase is very low compared to that of type I H^+^-PPase. Type II H^+^-PPases represent approximately 0.1 and 0.2% of the total H+-PPases in microsomal fractions of suspension cells and young roots, respectively (Segami et al., 2010). Beside that, the PPi hydrolysis activity wasn’t detected in *fugu5* mutants, indicating that the contribution of type II H^+^-PPase in cytosolic PPi hydrolysis is fairly insignificant (Ferjani et al., 2011).

The expression patterns and/or the protein accumulation of the type I H^+^-PPase suggested that it may play a role in accumulation of sugars and organic acids in vacuoles of peach (Etienne et al., 2002), pear (Suzuki et al., 1999) and grape (Venter et al., 2006). Type I H^+^-PPase was also suggested to play a role at early stages of fruit development at the cell division stage (Mohammed et al., 2012). The roles of Type I H^+^-PPase in growth and development are mediated by auxin transport (Li et al., 2005), or through the removal of excess levels of PPi to guarantee the forwardness of the sucrose-synthesis gluconeogenesis pathways (Ferjani et al., 2011). In contrast, information on the physiological roles of type II H^+^-PPases in plant growth and development is lacking.

Inorganic pyrophosphate (PPi) is a by-product of several biosynthetic pathways (e.g. Nucleic acids, proteins, lipids, and polysaccharide polymerization) (Stitt, 1998; Heinonen, 2001). PPi generating and consuming reactions appear to be close to equilibrium, thus it needs to be hydrolyzed to ensure the forwardness of the polymerization and PPi-producing reactions (Weiner, 1987).

There are at least three PPi-utilization enzymes in the cytosol of plant cell. (I) UGPase (UDP-glucose pyrophosphorylase) catalyzes the formation of Glucose-1-phosphate and UTP from UDPGlc and PPi during sucrose degradation by sucrose synthase (Weiner, 1987; Stitt, 1998). (II) PFP (Pyrophsophate-fructose 6-phosphate 1-phosphotransferase) which activates the phosphorylation of Fru6P using PPi as a phosphoryl donor (Weiner, 1987; Stitt, 1998). (III) The tonoplast-bounding proton pumping pyrophosphatase (H^+^-PPase) that energizes the secondary active transport utilizing the PPi as energy source (Weiner, 1987; Stitt, 1998).

PPi can serve as an alternative energy source during ATP deficiency (Stitt, 1998). In contrast to the strong decline in ATP level under hypoxic conditions, PPi level was found to be relatively invariable (Mustroph, 2005; Dancer, 1989; Geigenberger, 2000; Mustroph et al., 2005).

Tobacco and Potato plants with low PPi content due to the overexpression of *E. coli* inorganic pyrophosphatase, exhibited a strong defect in carbohydrate metabolism (Jelitto, 1992; Sonnewald, 1992). PPi removal affected the equilibrium between the reactants catalyzed by UGPase (Stitt, 1998). UGPase plays different roles in source and sink tissues. In sink tissues, it works in sucrose breakdown, while in source tissues it facilitates sucrose biosynthesis (Kleczkowski, 1994; Coleman et al., 2007).

This study is aiming to shed light on the physiological roles of H^+^-PPases in sugar starvation tolerance. To Do this, *fugu5,* a type I H^+^-PPase-loss of function mutant, and *avp2;1* and *avp2;2* mutants were used. Results suggest a possible role for H^+^-PPases in energy deprivation response possibly through influencing the phosphorylation activity of SnRK1 (Sucrose non fermenting protein kinase I), the central coordinator of transcriptional networks of the stress and energy signaling.

## Materials and methods

### Plant materials and growth conditions

*Arabidopsis thaliana* ecotype Columbia (Col-0) or mutants in Col-0 background were used in this study. Seeds of *fugu5* mutants, (*fugu5-1, fugu5-2, and fugu5-3*), were kindly provided by Dr. Ferjani (Ferjani et al., 2011). T-DNA insertion mutants *avp2;1* (SALK_054291), and *avp2;2* (SALK_138132C)) were obtained from The Arabidopsis Information Resource center (TAIR). Seeds were surface-sterilized by 500 μL of a sterilization solution, consists of 5% (v/v) sodium hypochlorite with 0.05% Tween 20 for 15 minutes. Seeds were then washed four times by sterilized distilled water. Plants were grown in half-strength MS medium only, or supplemented with 1% sucrose. All media were solidified by 0.7% agar and the pH was set to 5.7. Seeds were stratified at 4°C in the dark for 4 days, then they were incubated in growth chamber (NK system, Osaka, Japan) at 22°C under continuous light of 30 ~ 50 μmol m^-2^ s^-1^.

### Screening of H^+^-PPase-defective T-DNA insertion mutants

Conventional PCR was performed to confirm the T-DNA insertion sites and to isolate homozygous lines. The left and right oligonucleotides for *AVP2;1* (At1g78920) were GTGATTGGTATCGCCATCCT and ACAAACGAAGCACCAAAACC, respectively. For *AVP2;2* (At1g16780), the left primer was TGTCTTGCAGCCAGAAAGTG, and the reverse primer was TACCATTATGCCAGGCCTCT. The LBb1.3 T-DNA-specific primer was used in both cases.

### Expression analyses of *AVP*_and *DIN* genes

Total RNA was extracted by using RNeasy Plant Mini Kit (Qiagen, Hilden, Germany), following the manufacturer’s instructions. Genomic DNA contamination was eliminated by performing on-column digestion using RNase-free DNase I. RNA concentrations were measured by NanoDrop 1000 spectrophotometer (Thermo Fisher Scientific, Wilmington, USA). Complementary DNA (cDNA) was synthesized using Rever Tra Ace® qPCR RT Kit (Toyobo, Osaka, japan), following the manufacturer’s instructions. Quantitative Real Time PCR was performed with MyiQ PCR machine (Bio-Rad, Hercules, CA, USA), using SsoAdvanced SYBR® green supermix (Bio-Rad, Hercules, CA, USA). Gene expression results were normalized against *ACTIN2* gene expression.

List of primers used for *AVPs* and *DINs* expression analysis

AVP1F1790 : 5’CTCCTTGGCTCTCTTTGGTG3’
AVP1R1940 : 5’AGCTGCACTTCCCACACTCT3’
AVP2;1F2287 : 5’ATCCCAGAAGTGTTCGTTGG3’
AVP2;1R2473 : 5’CTATGGCGACACAGCGACTA3’
AVP2;2F2279 : 5’TTGTCGCATCTTCAGCTTTG3’
AVP2;2R2487 : 5’CCAAGCACCTCCTGCTGTAT3’
AtAct2+494F : 5’TGCCAATCTACGAGGGTTTC3’
AtAct2+719R : 5’TTCTCGATGGAAGAGCTGGT3’
DIN1F467 : 5’CAGAGTCGGATCAGGAATGG 3’
DIN1R572 :5’ATTTGACCGCTCTCACAACC 3’
DIN6F1301:5’ACTTGTCGCAAGATCAAG 3’
DIN6R1390 :5’GGAACACGTGCCTCTAGTCC 3’

### Extraction of total soluble proteins

Total soluble proteins for SnRK1 assay were extracted following the method of Zhang et al. (2009) with a few modifications. In brief, about 200 mg of seedlings grown in ½ MS medium were finely ground in liquid nitrogen using a pestle and a mortar. Ground tissues were re-suspended in 600 μL icecold homogenization buffer consists of 100 mM Tricine-NaOH, pH 8, 25 mM NaF, 5 mM dithiothreitol, 2 mM tetrasodium pyrophosphate, 0.5 mM EDTA, 0.5 mM EGTA, 1 mM bezamidine, 1 mM phenylmethylsulfonyl fluoride, Protease inhibitor cocktail (cOmplete; Roche Basel, Switzerland), Phosphatase inhibitors (PhosStop; Roche, Basel, Switzerland), and 2% (w/v) polyvinylpyrrolidone. Homogenates were centrifuged at 13000g for 15 minutes at 4°C. Supernatants were desalted with Illustra NAP-5 columns (GE Healthcare Life Sciences, Little Chalfont, UK) pre-equilibrated by homogenization buffer. Protease inhibitor cocktails and 2.5 μM Okadaic acid (Wako, Osaka, Japan) were added to the eluents before freezing in liquid nitrogen and storing at −80°C.

### SnRK1 activity assay

SnRK1 assay was performed following previously described protocols (Davies et al., 1989; Weekes et al., 1993; Dale et al., 1995; Zhang et al., 2009) with a few modifications. SnRK1 assay medium consists of 40 mM HEPES-NaOH, pH 7.5, 5 mM MgCl_2_, 100 μM ATP containing 12.5 kBq (γ-^33^P) ATP (PerkinElmer, Waltham, MA, USA), 200 μM AMARA peptide (H-Ala-Met-ALa-Arg-Ala-Ala-Ser-Ala-Ala-Ala-Leu-ALa-Arg-Arg-Arg-OH (Enzo® Life Sciences, Farmingdale, USA)), 5 mM dithiothreitol, 1 μM Okadaic acid, and protease inhibitor. Assay was started by adding 15 μL of the reaction mixture to 10 μL of the extracted total soluble protein. Incubated at 30°C for 6 minutes, 10 μL of the extract-reaction mixture was added to 1 cm^2^ P81 Phosphocellulose paper squares (MERCK MILLIPORE, Billerica, MA, USA). The papers were immersed directly in 1% (w/v) phosphoric acid (WAKO, Osaka, Japan). Then, they were washed four times in 1 mL 1% (w/v) phosphoric acid, immersed in acetone for 15 minutes, air-dried and then added to 3 mL of scintillation cocktail (Ultima Gold™; PerkinElmer, Waltham, MA, USA). Incorporated P^33^ in AMARA peptide was quantified using a liquid scintillation counter (Beckman, CA, USA).

### Determination of PPi level

PPi extraction was performed following the method of Kabala et al. (2010) with a few modifications. In brief, 200 mg of seedlings were ground in liquid nitrogen before suspending in 1.5 mL 5.4% (v/v) perchloric acid. The mixture was supplemented with 125 μL 2 mM Tris, and centrifuged at 7000g for 5 minutes. The pH of the supernatants was set to 7.4~7.6 by adding few drops of 5 M K_2_CO_3_, and re-centrifuged at 5000g for 5 minutes. The clear supernatant was used for measuring PPi concentration by using PiPer Pyrophosphate Assay Kit (Life Technologies, Carlsbad, CA, USA).

## Results

To understand the physiological function of H^+^-PPases in plant development, I used three lines of *A. thaliana* with defect in type I H^+^-PPase, namely *fugu5-1, fugu5-2,* and *fugu5-3* (Ferjani et al., 2011) and two homozygous T-DNA insertion lines, *avp2;1* and *avp2;2*, were isolated from Salk collection using gene-specific and T-DNA specific primers (Fig. 1). The expressions of both *AVP2;1* and *AVP2;2* in *avp2;1* and *avp2;2* mutants were reduced significantly (Fig. 2)

**Fig. 1.**
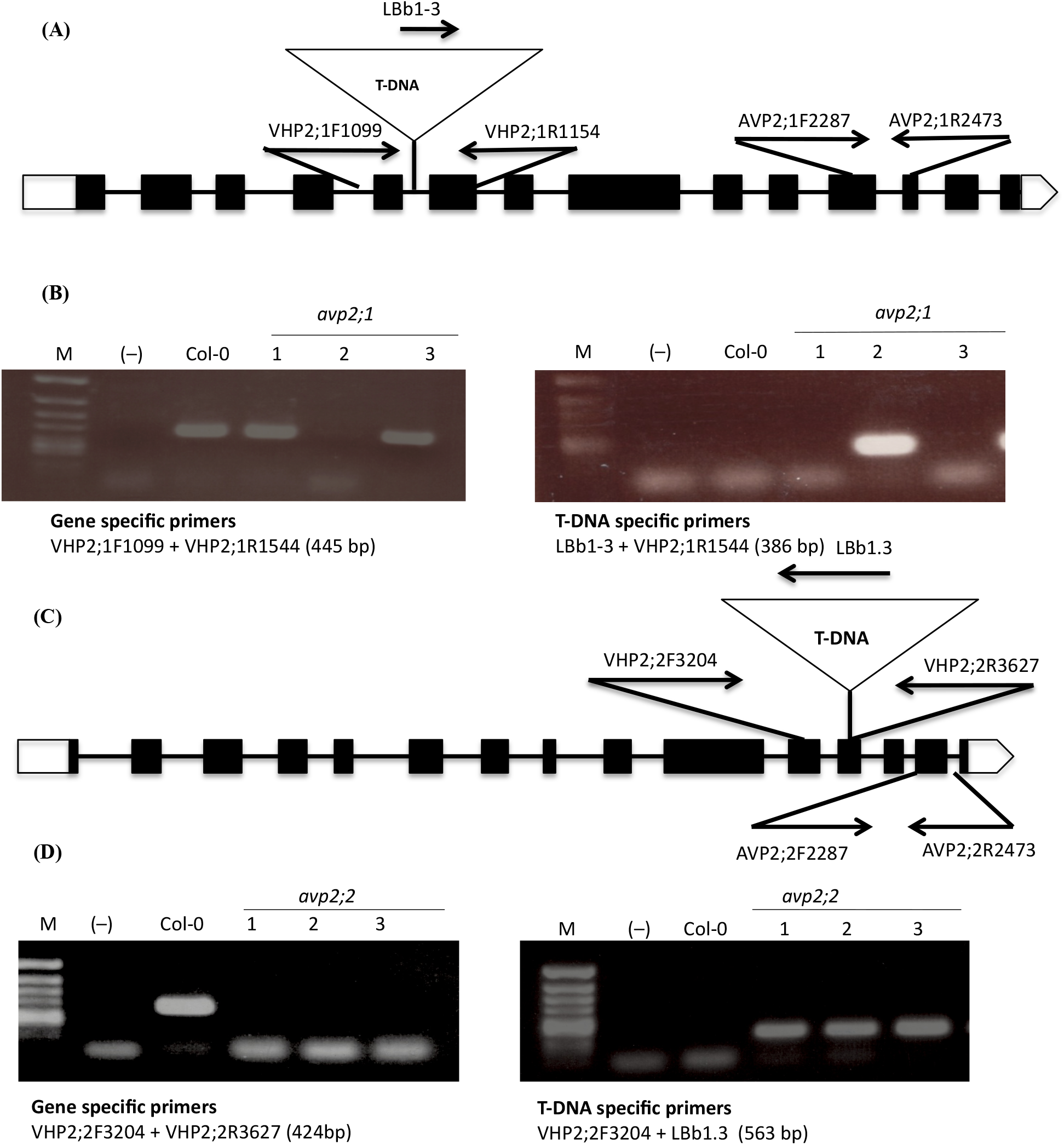
Isolation of avp2;1 *and avp2;2* mutants. Schematic diagrams show the position of T-DNA insertion in *avp2;1* **(A)** and *avp2;2* **(C).** Boxes indicate exons. Black boxes and white boxes indicate the translated and untranslated regions, respectively. Lines indicate introns. Small arrows represent primers, and the triangles indicate the T-DNA insertion. **(B)** and **(D)** Agarose gel electrophoresis of the amplified PCR products of WT and T-DNA insertion mutants. M lane indicates the *ϕX174 Hinc* II digest molecular marker, (-) is for negative control to which DNA template was not added. Col-0 is the wild type, and lane 1 to 3 are the examined plants of T-DNA insertion. lines No. 2 in (B), and No. 1 and 3 in (D) were homozygous mutations of T-DNA insertion.

**Fig. 2.**
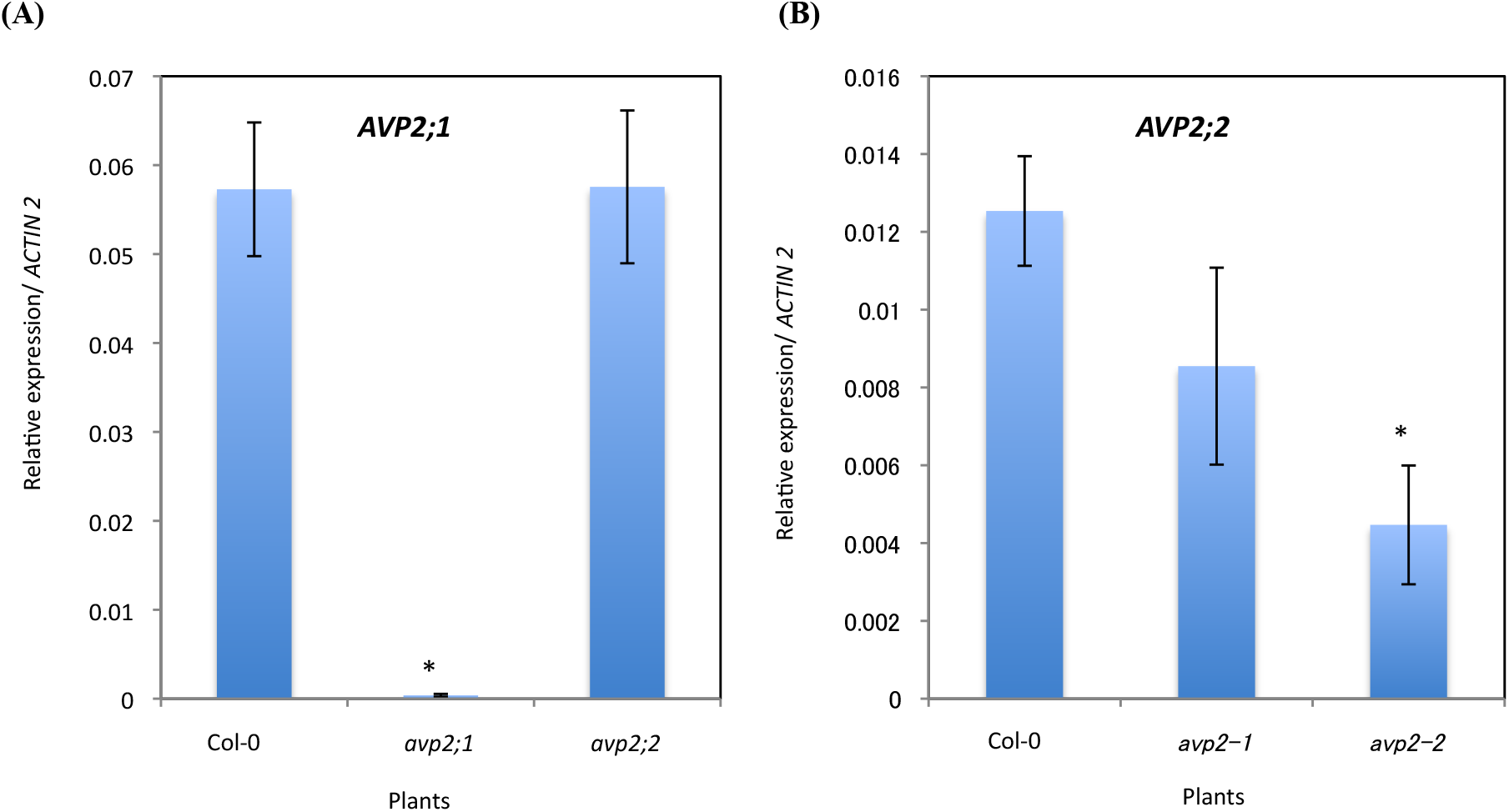
The reduced expression of *AVP2;1,* and *AVP2;2* in isolated type II H+-PPase-defective mutants. **(A)** *AVP2;1* gene expression. **(B)** *AVP2;2* gene expression. Each datum is the mean of three independent experiments, and represented as means ± SE. Asterisks indicate a value that significantly different from that of Co-0 at *P* < 0.05 using student’s *t*-test.

### *fugu5* mutants accumulate high level of PPi

To determine the level of intracellular PPi, the soluble metabolites were extracted in 5.4% perchloric acid, and then the PPi concentration was measured through formation of the fluorescent resorufin by using PiPer Pyrophosphate assay kit. The results indicated that all *fugu5* mutants accumulate significantly higher level of PPi than the wild-type plants (Fig. 1). The PPi levels in *avp2;1* and *avp2;2* mutants were significantly indifferent from that of wild-type plants (Fig. 3), suggesting that the role of type II H^+^-PPase in PPi hydrolysis could be minor.

**Fig. 3.**
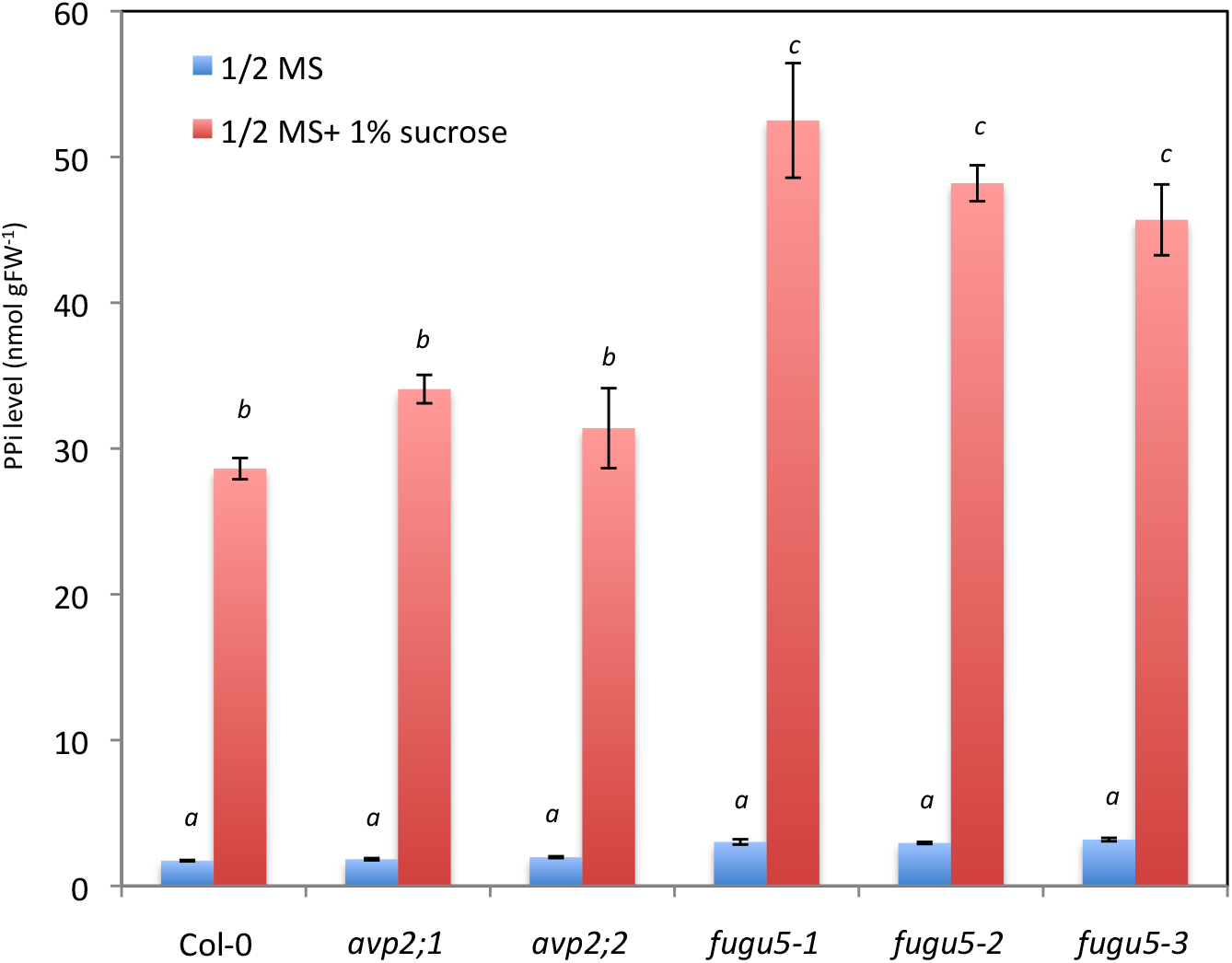
PPi accumulation in *fugu5* mutants. Plants were grown for one week on sugar-free ½ MS medium, or on ½ MS supplemented with 1% sucrose. Data represent the mean ± SE for three replicates. Different letters indicate statistically significant difference at *P* < 0.05 by using Tukey’s HSD method.

### *fugu5* mutants showed higher tolerance to sugar starvation

As a way of coping with energy deprivation, plants show a growth arrest and redirection of all cellular activity toward basic metabolism (Rolland et al., 2006). It is known that PPi can serve as an alternative energy source during ATP deficiency (Stitt, 1998). Because *fugu5* mutants accumulate significantly higher level of inorganic pyrophosphate PPi, resulting from to the loss of function of H^+^-PPase, it was interesting to investigate the effect of energy deprivation on the growth of the mutants. The wild type, *avp2;1, avp2;2, fugu5-1, fugu5-2,* and *fugu5-3* plants were grown on sugar-free half-strength MS medium and incubated at 22°C under low light intensity of 30 ~ 50 μmol m^-2^ S^-1^. Under these photosynthesis-constrained conditions, the wild-type plants exhibited growth and developmental arrest. This is mainly because of the low-light intensity, which wasn’t enough to support the full autotrophic growth of seedlings when cultured on sugar-free medium. In contrast, the *fugu5* outperformed the wild type and developed longer root systems (Fig. 4 and Fig. 5).

**Fig. 4.**
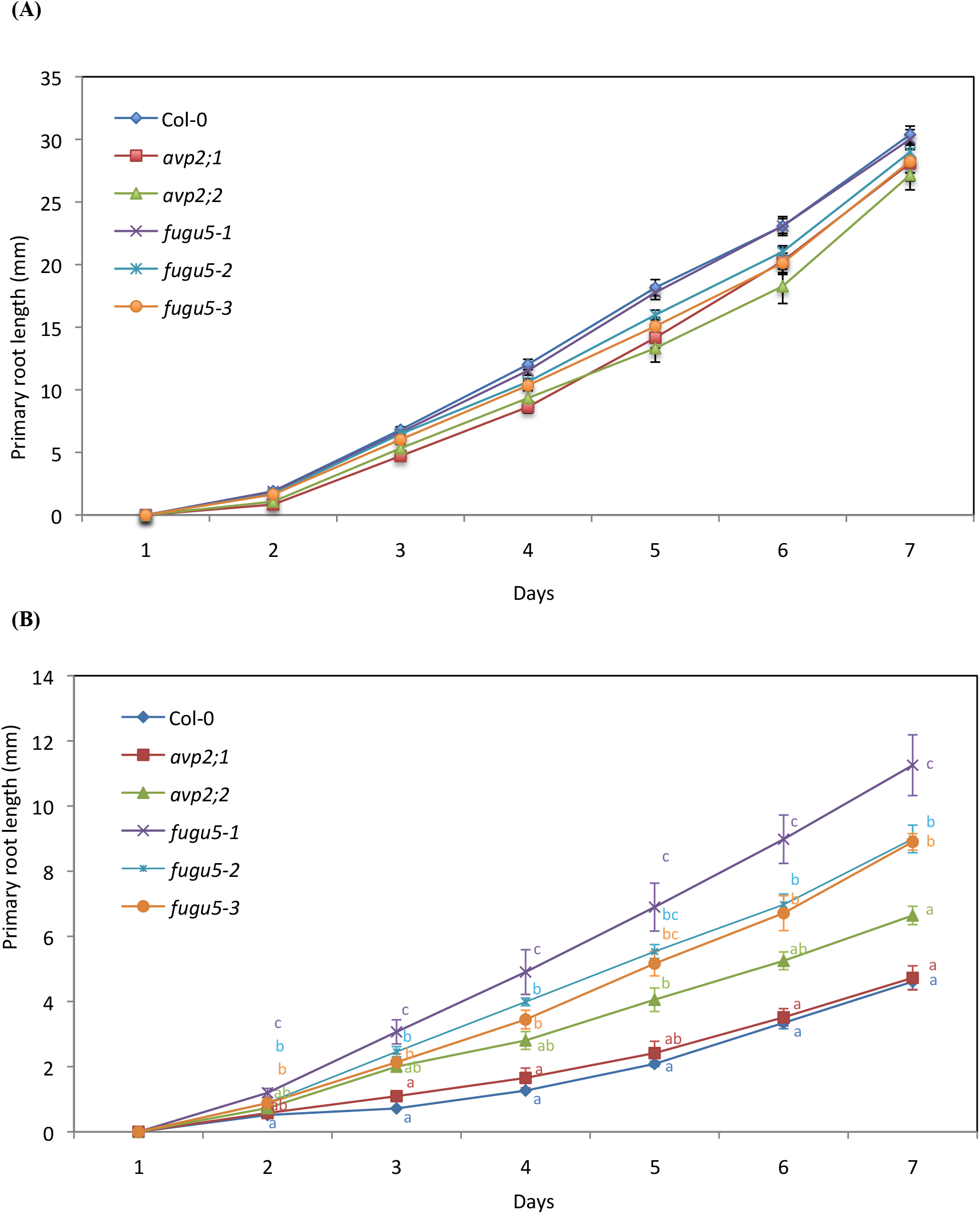
Effect of sugar starvation on primary root growth of H^+^-PPase-defective mutants. Plants were grown on sugar-free ½ MS medium **(A)**, or on ½ MS supplemented with 1% sucrose (B).The root growth was monitored daily, and the root length was measured using ImageJ software. All plants were grown under low light conditions. Data represent the mean ± SE. 15 ≤ *n* ≤ 20 for (A) and 30 ≤ *n* ≤ 60 for **(B)**. Different letters indicate statistically significant difference at *P* < 0.05 by using Tukey’s HSD method.

**Fig. 5.**
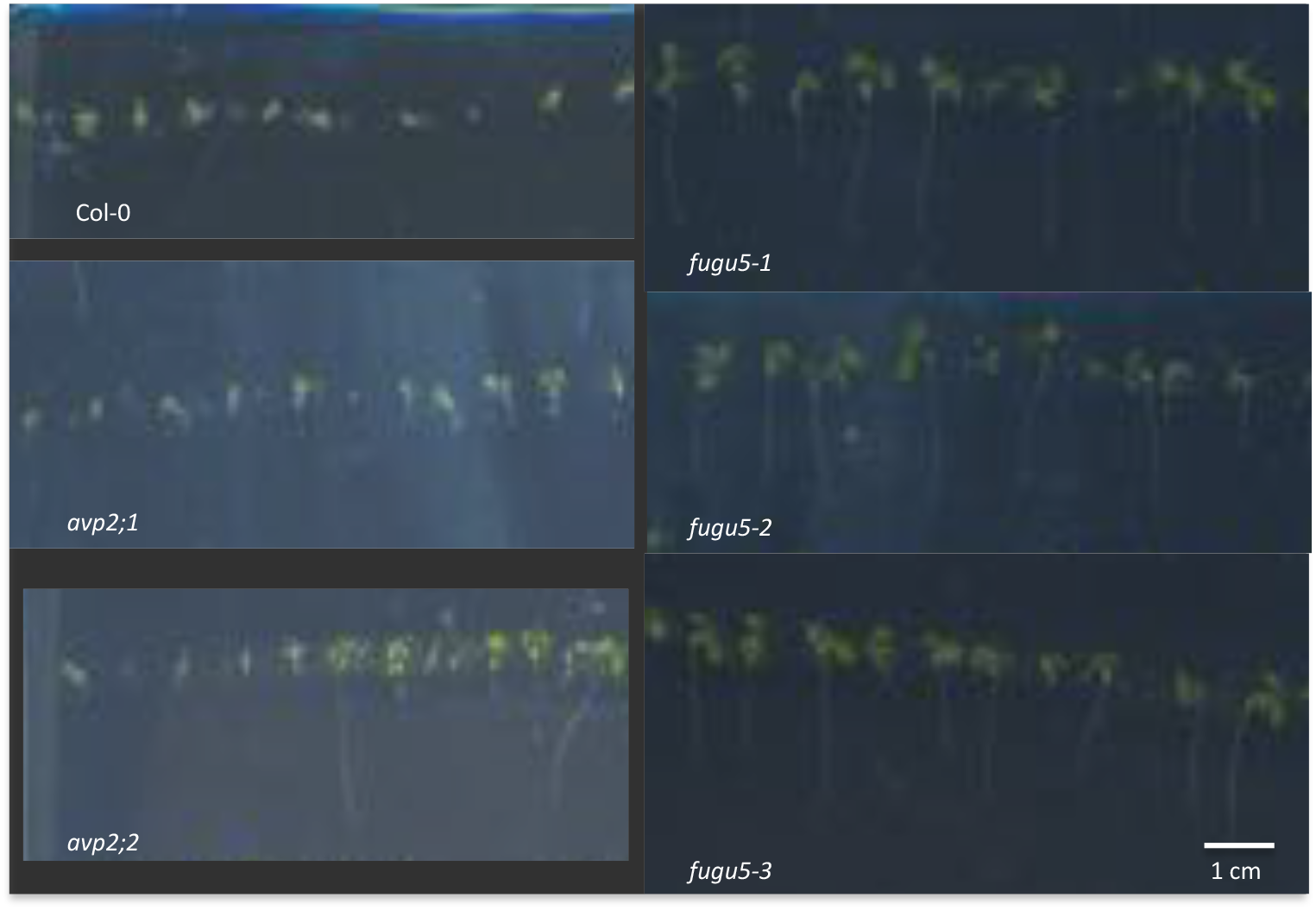
Effect of sugar starvation on primary root growth of H^+^-PPase-defective mutants. Plants were grown for 7 days on sugar-free ½ MS medium, under low light condition.

### The expression of the genes encoding H^+^-PPase is reduced under sugar starvation conditions

Sugar regulates the expression of many genes directly or through interacting with many other signaling networks (Morkunas et al., 2012). To gain more insight into the role of H^+^-PPase in sugar starvation, I compared the expression level of *AVP1, AVP2;1,* and *AVP2;2* genes in wild-type seedlings subjected to sugar starvation, with their expression in 1% sucrose-containing ½ MS medium. Results indicated a remarkable reduction in expression of all genes encoding H^+^-PPase. *AVP1* expression was reduced by 74 %, *AVP2;1* by 79%, and *AVP2;2* by 76 % (Fig. 6).

**Fig. 6.**
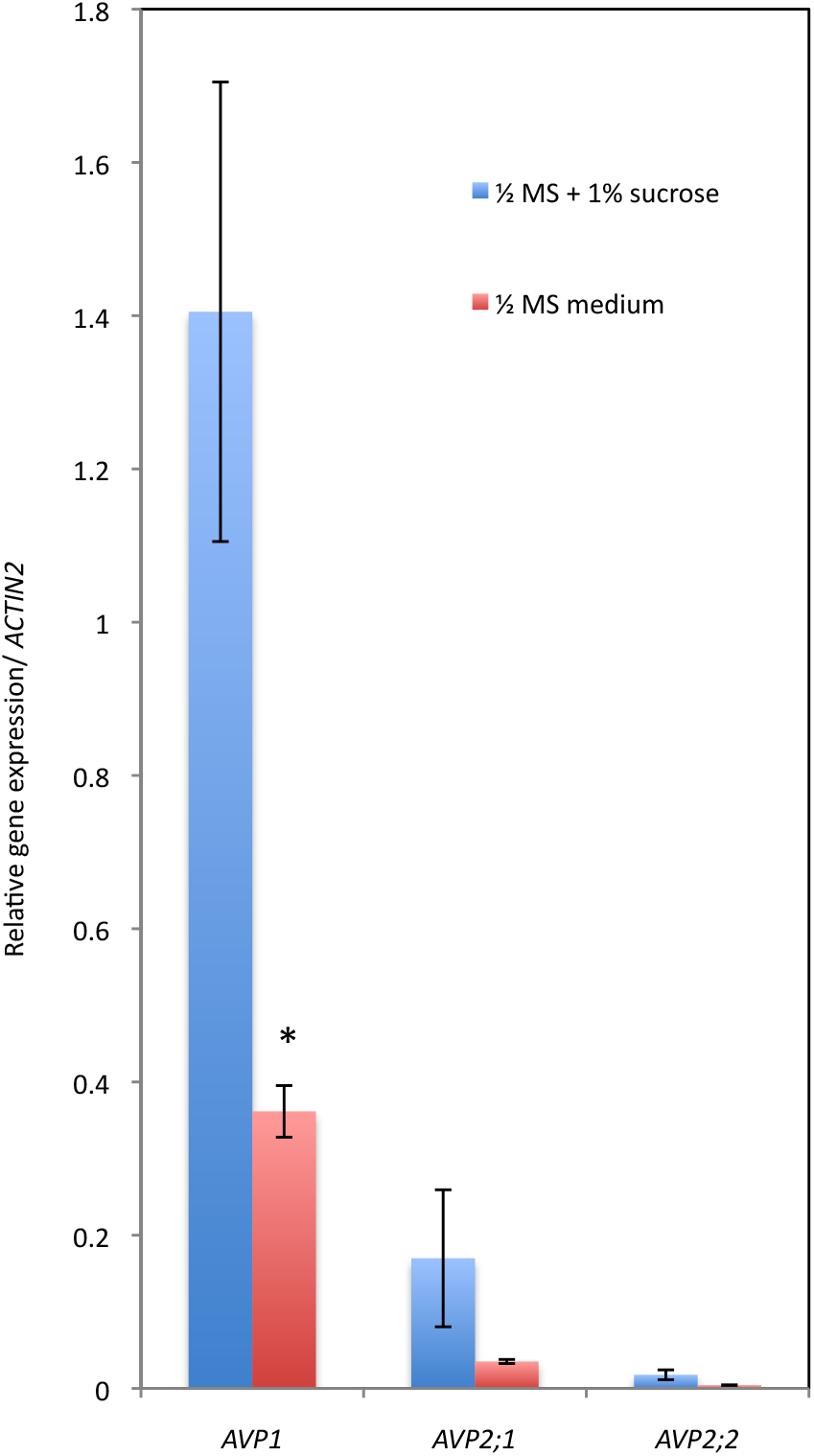
Effect of sugar starvation on H^+^-PPase-encoding genes expressions. RNA was extracted from plants grown for one week on ½ MS medium under low light condition. Data represent mean ± SE of three replicates. Gene expression was normalized against *ACTIN2* gene. Asterisk indicates a value that significantly different from that of Co-0 at *P* < 0.05 using student’s *t*-test.

### *DIN1* and *DIN6* gene expression under sugar starvation conditions

Dark-Inducible genes (*DINs*) are transcriptionally activated by energy depletion in darkness, sugar starvation and stress conditions (Baena-González et al., 2007). The tolerance response of *fugu5* mutants to sugar starvation was higher than that of wild-type plants; therefore, I investigated the expression of *DIN1* (At4g35770) and *DIN6* (At3g47340) in H^+^-PPase-defective mutants. *DIN1* encodes for a senescence-associated protein, and functions in RNA regulation. *DIN6* encodes for a glutamine-dependent asparagine synthetase and functions in amino acid metabolism. Both gene expressions are known to be induced in the dark and under sugar starvation. Results showed that, the level of *DIN1* transcripts in *avp2;1, avp2;2, fugu5-1, fugu5-2,* and *fugu5-3*, in sugar starvation medium under light condition were 34%, 64%, 45%, 53%, and 48% lower than that of control plants, respectively (Fig. 7 A). The induction of *DIN6* by sugar starvation was remarkably reduced in H^+^-PPase-defective mutants compared to wild-type plants (Fig. 7 B). These results indicate that under low-light conditions, the induction of *DINs* genes by sugar starvation in H^+^-PPase-defective mutants is lower than that of wildtype plants.

**Fig. 7.**
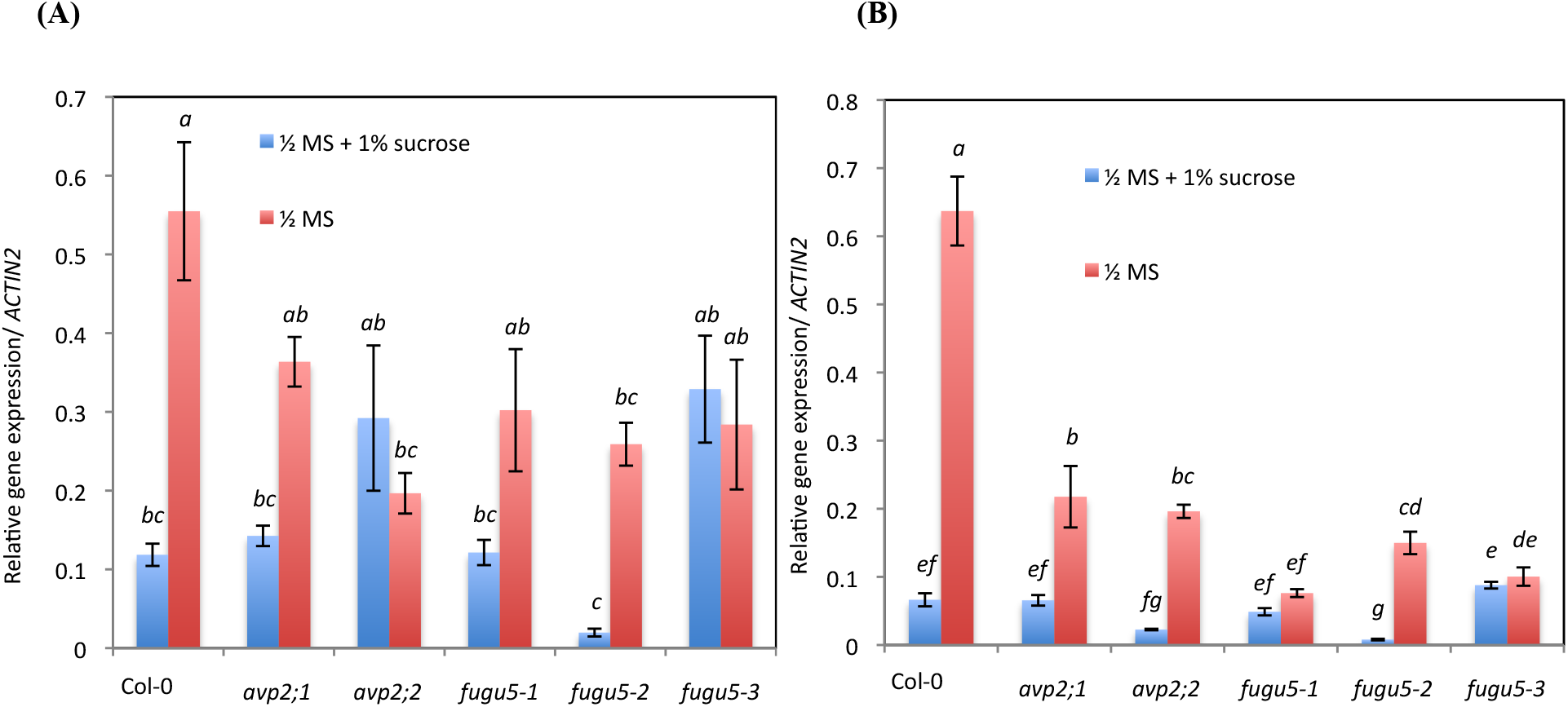
Effect of sugar starvation on *DIN1* and *DIN6* gene expression of H^+^-PPase-defective mutants. **(A)** *DIN1* gene expression. **(B)** *DIN6* gene expression. RNA was extracted from plants grown for one week on ½ MS medium under low light condition. Data represent mean ± SE of three replicates. Gene expression was normalized against *ACTIN2* gene.

### SnRK1 activity is reduced in *fugu5* mutants

To further understand the mechanisms that caused the better growth performance of H^+^-PPase mutants, I assayed the phosphorylation activity of SnRK1. SnRK1 is functionally and structurally related to the yeast sucrose-non-fermenting-1-kinase and animals AMP-activated protein kinase (Hey et al., 2007). SnRK1 senses the metabolic status and allow plants to adapt to stress conditions by promoting catabolism and inhibiting anabolism, and thus it saves ATP to be used for pivotal cellular processes (Tome et al., 2014). AMARA peptide, the specific substrate for SnRK1 (Dale et al., 1995; Salt et al., 1998) and radiolabeled ATP have often been used to investigate the SnRK1 activity. The SnRK1 phosphorylation activity in extracts of total soluble proteins of seedlings was induced significantly by sugar starvation in wild type, *avp2;1* and *avp2;2* mutants, but not in *fugu5* mutants. The reduction of SnRK1 activity could be one of the reasons behind the better growth performance of *fugu5* mutants under sugar starvation conditions (Fig. 8).

**Fig. 8.**
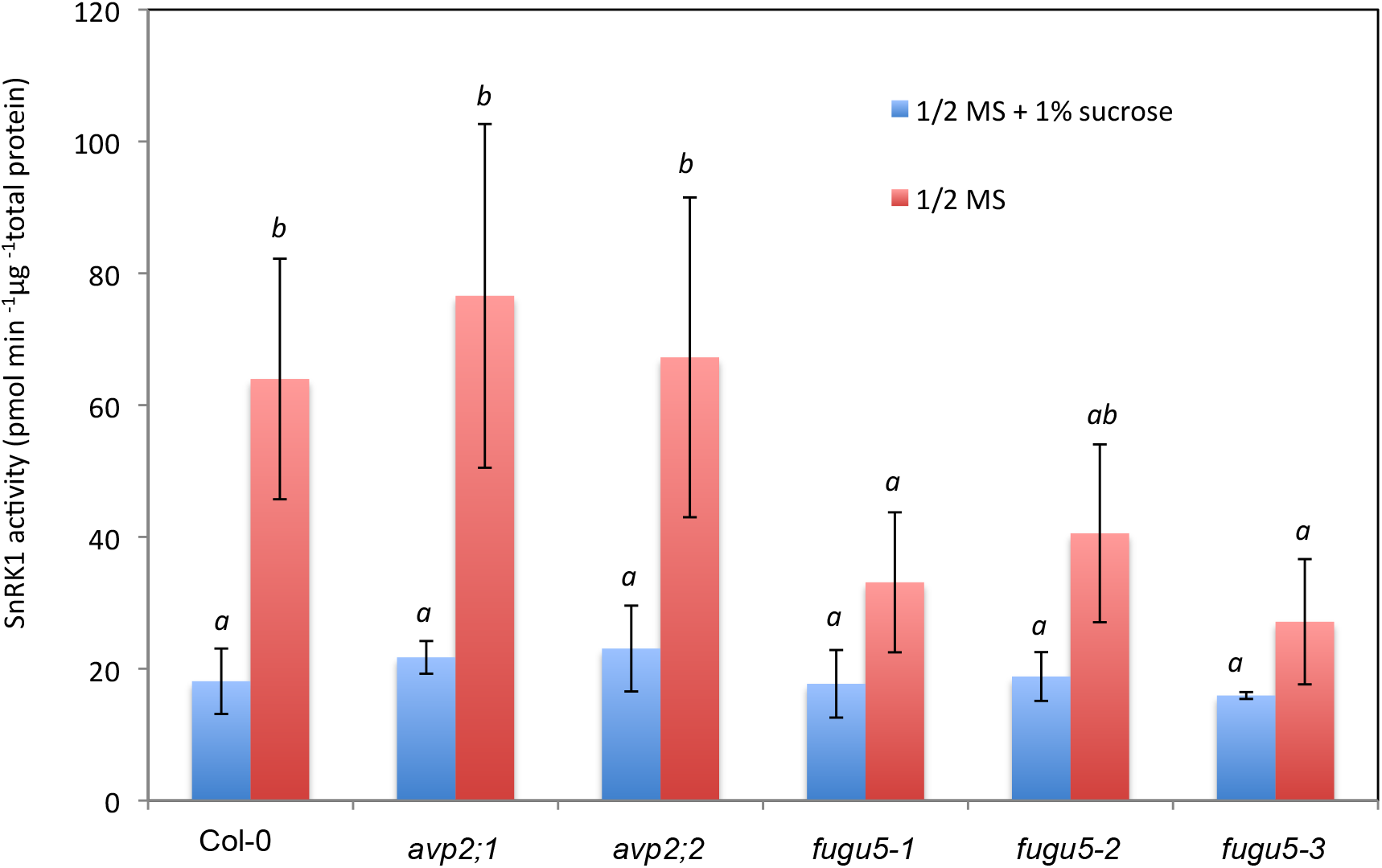
Reduction of SnRK1 activity in H^+^-PPase defective-mutants. SnRK1 activity was measured by using a liquid scintillation counter after removing unincorporated (γ^33^P) ATP from reaction mixture. Total protein was extracted from plants grown for one week on ½ MS medium under low light condition. Data represent mean ± SE of three replicates. Different letters indicate statistically significant difference at *P* < 0.05 by using Tukey’s HSD method.

## Discussion

Plants have a sessile nature. Therefore, they must continuously sense and coordinate their metabolism with internal and external signals to optimize growth and development (Tomé et al., 2014). Sugars, particularly sucrose and glucose, play crucial roles in this coordination. Sugars function not only as energy sources and structural components; they also work as central regulators for metabolism, physiology, development, and gene expression of many vital processes such as photosynthesis, respiration, cell cycle regulation, nitrogen metabolism, pathogen defense, wounding response, and senescence (Jang et al., 1997). Stress conditions such as nutrient scarcity, hypoxia, salts, and extreme temperatures; often affect photosynthesis and respiration in source leaves. Therefore, it can lead to growth arrest due to the massive alteration in cellular process (Tomé et al., 2014).

In this study, the wild-type plants exhibited growth arrest phenotype, when grown in sugar free basal medium under low light conditions. The low photosynthesis wasn’t able to support the full autotrophic growth of wild-type plants. Xiong et al. (2013) showed a similar phenotype in Arabidopsis plants grown in sugar-free medium and low light intensity of 75μmol m^-2^s^-1^. In this study Arabidopsis plants were grown at even weaker light intensity of 30 - 50 μmolm^-2^s^-1^. In general, plants react with sugar starvation and energy deprivation by arresting growth and redirecting cellular activities toward basic metabolism, by enhancing catabolism and inhibiting anabolism (Rolland et al., 2006).

Beside their roles as energy sources, sugars also regulate expression of hundreds of genes in plants (Kunz et al., 2014). It has been shown that the expression of 983 genes is upregulated, whereas the expression of 771 genes is downregulated in Arabidopsis in response to glucose at 3% (Li et al., 2006). The expression of vacuolar pyrophosphatase genes is downregulated during sugar starvation. This suggest that the inhibition of these genes spares their substrate, PPi, to be used for coping with starvation conditions.

The *fugu5-3* mutant accumulates 50% more PPi than wild-type plants indicating that *AVP1* is the main hydrolyser of the cytosolic PPi (Ferjani et al., 2011). I found that beside *fugu5-3, fugu5-2* and *fugu5-1* accumulate significantly higher levels of PPi. PPi producing and utilizing reactions are reversal. The high PPi level could inhibit PPi-producing reactions (e.g. polymerization reactions), and thus it was expected to affect plant growth negatively. Contrary to the expectations, however, all *fugu5* mutants displayed higher tolerance to sugar starvation and better performance in term of growth than wild-type plants.

Results propose that the phenotype of starvation tolerance could be attributed, at least in part, to the high amount of PPi accumulated in the cytosol of *fugu5* mutants. High nitrogen level in growth medium enhances plant growth. Adequate sucrose concentration is necessary for this enhancement (George et al., 2007). This is possibly one of the main reasons of severe growth arrest in full strength compared to half strength MS medium under low light conditions (Data not shown).

Energy depletion can affect the expression of many genes (Usadel et al., 2008). These genes form a network that regulates plant metabolism under stress conditions for optimum energy preservation (Avin-Wittenberg et al., 2012). The expression level of H^+^-PPase-encoding genes was reduced significantly (Fig. 5). It is possibly a way of energy conservation through increasing of PPi level, which can replace ATP for powering some reactions (Stitt, 1998).

*DIN* genes expression was reduced in H^+^-PPase-defective mutants comparing to wild-type plants. However, the expression was not correlated with the growth-retarded phenotype of *fugu5* mutants, as both type I and II H^+^-PPase-defective mutants showed a reduced expression for *DINs. DIN* genes work downstream of SnRK1. The phosphorylation activity of SnRK1 indicates the energy status of plants.

Interestingly, the transgenic Arabidopsis plants overexpressing *OsSnRK1* showed hypersensitivity to high glucose concentrations, while the transgenic plants expressing inactive *OsSnRK1* were insensitive (Cho et al., 2012). SnRK1 is a master regulator and central coordinator of plant transcription networks to stress and energy signaling (Baena-González et al., 2007). Transient elevation of SnRK1 results in activation of many genes involve positively in stress- and starvation-responses, or inhibition of expressions of many genes correlated with activation by sucrose, glucose, and elevated CO2 (Sheen, 2014). I examined the SnRK1 activity in H^+^-PPase-defective mutants. The results showed that the activity was reduced significantly in *fugu5* mutants compared to wild-type and type II H^+^-PPase mutant plants under sugar-starvation conditions. This result suggests that the high PPi level in *fugu5* mutants is responsible, at least in part, for the reduced activity of SnRK1.

ATP available for different cellular process could mediate this reduction. PPi can serve as an alternative energy source during ATP deficiency (Stitt, 1998). In contrast to the strong decline in ATP level under hypoxic conditions, PPi level was found to be relatively invariable (Dancer & Rees, 1989; Geigenberger et al., 2000; Mustroph et al., 2005). The high PPi level might help in conservation of ATP, probably through activation of PPi-dependent enzymes such as sucrose synthase, UGPase, and PEP rather than the ATP-depending enzymes. The conserved availability of ATP reduces the starvation, and this may explain the reduced SnRK1 activity. I propose a hypothetical model for the PPi-mediated sugar starvation tolerance (Fig. 9). Plants on MS medium rely partially on exogenous sugar till the photosynthetic apparatus becomes competent (Leon and Sheen, 2003). The low light intensity isn’t enough to support the full autotrophic growth (Xiong et al., 2013). The low light and the absence of sugars from the growth medium induce energy depletion. The low ATP:AMP ratio due to the starvation activates SnRK1 by phosphorylation of the catalytic subunit of the complex (Mohannath et al., 2014). Phosphorylated SnRK1 will control the plant transcriptome by phosphorylation of several transcription factors controlling genes that promote catabolic reactions and inhibit anabolic reactions (Tome et al., 2014). The growth is arrested as a result of regulation by SnRK1. When sucrose is available in the nutrient medium, the high ATP:AMP ratio will reduce the SnRK1 activity. This in turn will ease the control over growth and anabolic reactions. The results suggest that the high PPi level in *fugu5* mutants might contribute to conservation of ATP level. Sucrose degradation by invertase followed by phosphorylation of resulted hexoses by fructokinase and hexokinase requires two molecules of ATP. Whereas, sucrose degradation by sucrose synthase (SUSY) followed by phosphorylation of fructose by fructokinase requires only one molecuole of ATP with UGPase requiring one molecule of PPi. The formed UTP can be converted to ATP by nucleoside-5-diphosphate kinase and used to phosphorylate fructose (Dancer et al., 2005). Fructose-6-phosphate is phosphorylated by ATP-dependent phosphofructokinase (PFK) or PPi-dependent phosphofructokinase. Results suggest that the PPi-dependent pathways have the preference over the ATP-dependent pathways during sugar starvation under low light conditions in Arabidopsis.

**Fig. 9.**
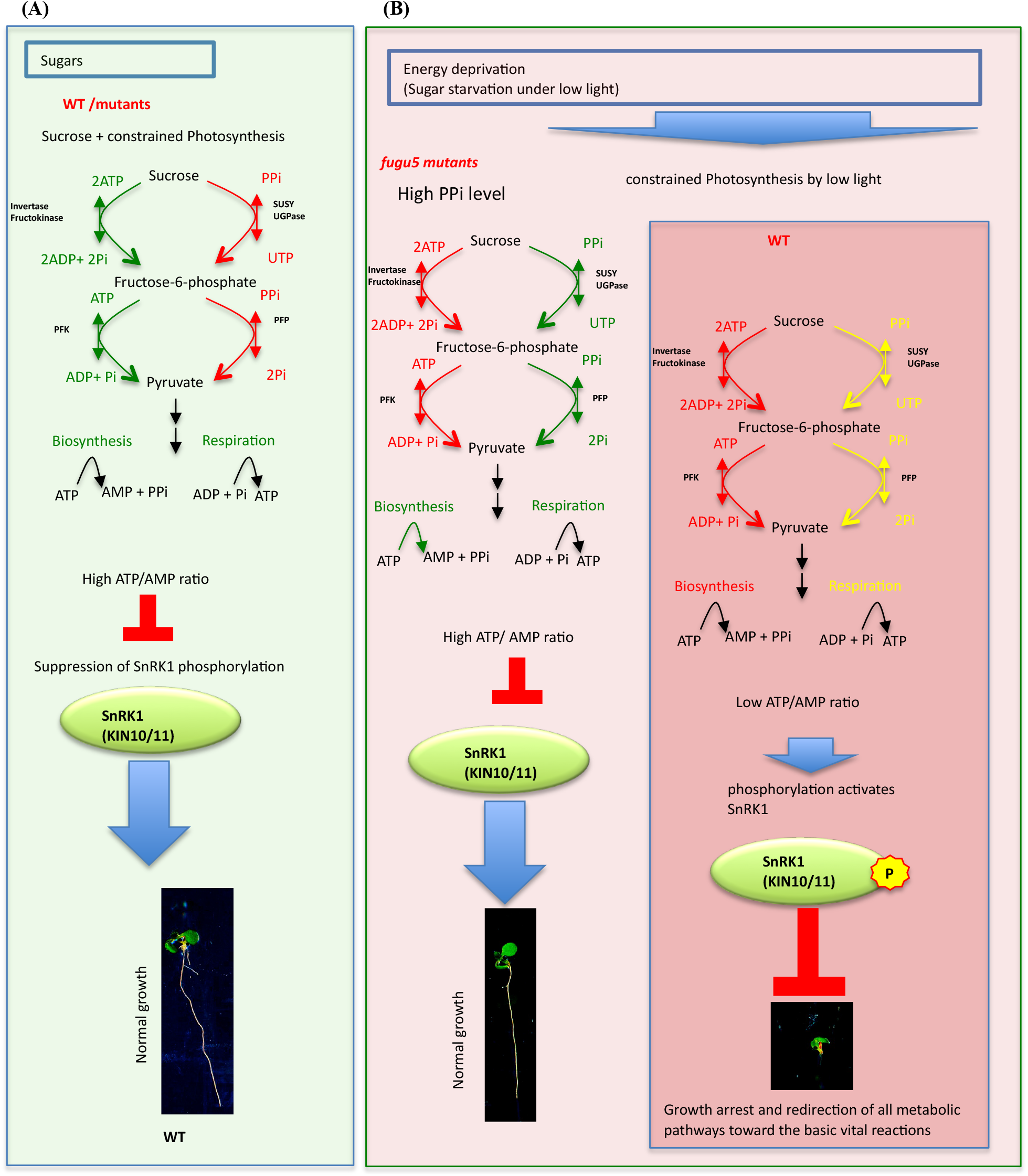
A hypothetical model for PPi-mediated sugar starvation tolerance. Sucrose degradation by invertase followed by phosphorylation of resulted glucose and fructose by frucktokinase requires two molecules of ATP. Sucrose degradation by sucrose synthase (SUSY) followed by UGPase require only one molecule of PPi. Fructose-6-phosphate is phosphorylated by ATP-dependent phosphofructokinase (PFK), or PPi-dependent phosphofructokinase. **(A)** Sucrose in growth medium is sufficient to support the semi-heterotrophic growth under low light condition. The ATP dependent reactions have the advantage over the PPi dependent reactions. High ATP to AMP ratio inhibits Sucrose-non-fermenting related protein kinase 1 (SnRK1) activity, which controls gene expression and protein phosphorylation in a way that induces catabolism and inhibits anabolism. **(B)** The low light constrains photosynthesis, and the absence of sugar from the growth medium induces sugar starvation and arrests plant growth in wild-type plants. The low ATP/AMP ratio activates SnRK1 through phosphorylation. The high PPi level in H^+^-PPase-defective mutants facilitates PPi-dependent reactions and conserve ATP. The comparatively high ATP/AMP ratio inhibits SnRK1.

